# Benchmarking a foundational cell model for post-perturbation RNAseq prediction

**DOI:** 10.1101/2024.09.30.615843

**Authors:** Gerold Csendes, Kristóf Z. Szalay, Bence Szalai

## Abstract

Accurately predicting cellular responses to perturbations is essential for understanding cell behaviour in both healthy and diseased states. While perturbation data is ideal for building such predictive models, it is considerably sparser than baseline (non-perturbed) cellular data. To address this limitation, several foundational cell models have been developed using large-scale single-cell gene expression data. These models are fine-tuned after pre-training for specific tasks, such as predicting post-perturbation gene expression profiles, and are considered state-of-the-art for these problems. However, proper benchmarking of these models remains an unsolved challenge.

In this study, we benchmarked a recently published foundational model, scGPT, against baseline models. Surprisingly, we found that even the simplest baseline model - taking the mean of training examples - outperformed scGPT. Furthermore, machine learning models that incorporate biologically meaningful features outperformed scGPT by a large margin. Additionally, we identified that the current Perturb-Seq benchmark datasets exhibit low perturbation-specific variance, making them suboptimal for evaluating such models.

Our results highlight important limitations in current benchmarking approaches and provide insights into more effectively evaluating post-perturbation gene expression prediction models.

## Introduction

Modelling cellular phenotypes is a fundamental challenge in computational systems biology. Accurately predicting cell fate can advance our understanding of both healthy and diseased states, and facilitate the identification of novel therapeutic targets. Over the past few decades, computational models based on Boolean logic (Montagud et al. 2022) and ordinary differential equations (ODEs) have been developed (Molinelli et al. 2013). More recently, advances in deep learning methodologies have reinvigorated interest in this area (Yuan et al. 2021; Theodoris et al. 2023; Roohani, Huang, and Leskovec 2023; Cui et al. 2024; Hao et al. 2024; Bunne et al. 2024).

Cellular phenotypes can be described using various data modalities, including transcriptomics, (phospho)proteomics, and imaging-based phenotypic assays (phenomics). Among these, transcriptomics data - derived from techniques such as microarrays, bulk RNA sequencing, and single-cell RNA sequencing (scRNA-seq) - is the most used for large-scale cellular phenotype analysis due to its relatively low cost and well-established analysis methods. While gene expression profiles do not directly reflect protein-mediated signalling, they offer a proxy for the overall cellular state (Dugourd and Saez-Rodriguez 2019). Notably, post-perturbation transcriptomics data are particularly suited for training computational models because the causal relationship between known perturbations (cause) and the measured post-perturbation gene expression (effect) allows for the modelling of mechanistic processes (Lamb et al. 2006). However, acquiring such perturbation data is more complex than obtaining baseline (non-perturbed) transcriptomics data.

To address these challenges, several Transformer-based, foundational cell models have emerged recently (Theodoris et al. 2023; Cui et al. 2024; Hao et al. 2024). These models are pre-trained on vast amounts (>10M examples) of unlabeled scRNA-seq data, with the aim that such large datasets allow the models to capture general principles of gene regulation and signalling. These pre-trained models can then be fine-tuned on perturbation data to predict post-perturbation phenotypes more effectively.

These models had shown strong performance in post-perturbation RNA-seq prediction tasks using Perturb-seq-based genetic perturbation benchmarks (Dixit et al. 2016). These models generally predict the post-perturbation RNA-seq profiles of perturbed single cells by using gene expression data from unperturbed cells, along with a representation of the perturbation. The primary evaluation for these models is their ability to predict RNA-seq profiles for unseen perturbations.

However, benchmarking machine learning models in this domain is challenging. We have previously demonstrated that test set selection and performance metrics can significantly impact benchmarking outcomes in the related problem of post-perturbation viability prediction (Szalai et al. 2023). Poor test set design or inappropriate metric choice can result in indistinguishable performance between well-performing and trivial models. Furthermore, computational models of cellular phenotypes have diverse applications, such as predicting the effects of known perturbations in novel cell types (Cell Exclusive, CEX setup) or predicting the effects of novel perturbations in familiar cell types (Perturbation Exclusive, PEX setup). Unfortunately, current benchmarks, which predominantly rely on Perturb-seq datasets comprising diverse genetic perturbations in a single cell line, primarily assess PEX performance, limiting their ability to evaluate generalisation in a broader context.

In this study, we benchmarked a recent Transformer-based foundation model, scGPT, across three Perturb-seq datasets. We compared its performance to baseline models of varying complexity. Surprisingly, we found that scGPT generally underperformed compared to a simple baseline model that uses the mean of the training samples. Standard machine learning models using biological prior-knowledge outperformed scGPT by a large margin. We further identified that the low inter-sample variance in commonly used datasets complicates model performance assessment.

## Results

### Benchmarking of post-perturbation RNAseq prediction methods

scGPT is a masked language model (LLM) based transformer architecture, pre-trained on large-scale, unlabeled single-cell RNA sequencing (scRNA-seq) data. Through pre-training, scGPT learns gene embeddings and captures gene-gene relationships, which can then be leveraged for various downstream tasks, including post-perturbation RNA-seq prediction. The model takes as input RNA-seq vectors from randomly selected unperturbed cells, along with a perturbation representation, to predict RNA-seq profiles of perturbed cells. scGPT uses a perturbation token, which is added to the perturbed gene token to model perturbation effects.

For benchmarking, we employed the three datasets from the scGPT paper. These datasets were generated using Perturb-seq, which combines CRISPR-based perturbations with single-cell sequencing to capture post-perturbation gene expression profiles. Specifically, the Adamson (Adamson et al. 2016) dataset comprises 68,603 single cells subjected to single perturbation CRISPR interference (CRISPRi). The Norman dataset (Norman et al. 2019) includes 91,205 single cells subjected to single or dual CRISPRa (overexpression). Lastly, a subset (Roohani, Huang, and Leskovec 2023) of the Replogle (Replogle et al. 2022) dataset containing 162,751 single cells from a genome-wide single perturbation CRISPRi screen was used.

For evaluation, we adopted the metrics used by the scGPT authors (Figure 1A). Predictions were generated at the single-cell level, and the predicted gene expression profiles for each perturbation were averaged to form pseudo-bulk expression profiles. These predicted profiles were then compared to the ground truth pseudo-bulk profiles using Pearson correlation coefficients. Importantly, Pearson correlations were calculated not only in the raw gene expression space but also in the differential expression space (i.e., perturbed gene expression profile minus control gene expression profile). Additionally, we evaluated performance on the top 20 differentially expressed genes to emphasise the model’s ability to capture the most significant transcriptional changes.

**Figure 1.**
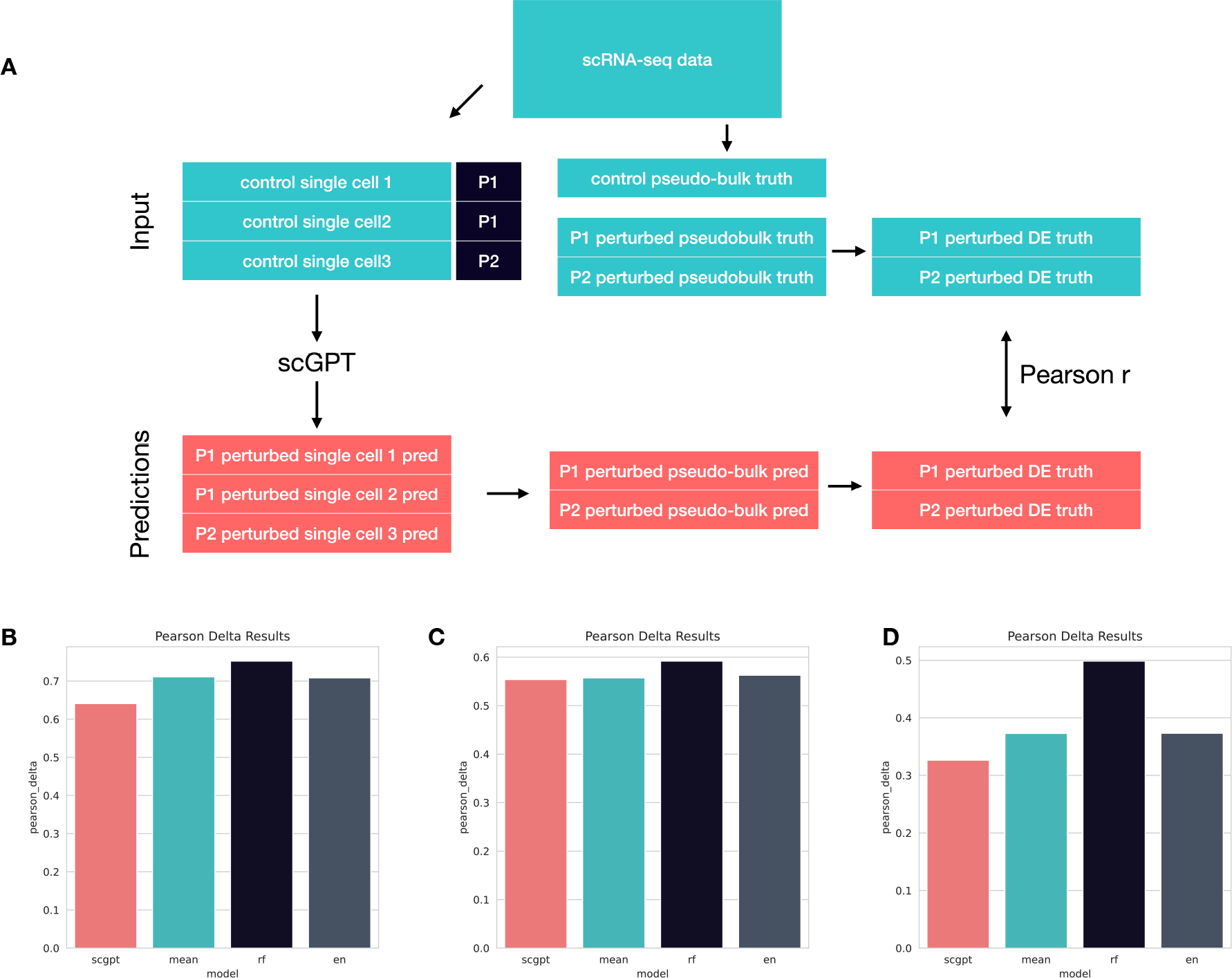
Benchmarking scGPT and baseline models. (A) Schematic representation of benchmark pipeline (B) Evaluation of Adamson dataset: Pearson delta metrics (y axis) for scGPT and 3 benchmark models (train mean, Random Forest Regression and Linear model) (x axis) (C) Evaluation of Norman dataset (D) Evaluation of Replogle dataset.

To further assess prediction performance, we introduced several baseline models. The simplest model, “Train mean”, predicted post-perturbation expression by averaging the pseudo-bulk expression profiles from the training dataset. We also constructed Linear Regression (LR) and Random Forest Regressor (RF) models. These models took as input the Gene Ontology (GO) vectors (The Gene Ontology Consortium 2019) of the perturbed genes. For the latter models, we used pseudo-bulked expression profiles on the target side (detailed in Methods).

Our results show that reproduced scGPT models reached similar performance to the original manuscript (Figure 1B, 1C, 1D; Table 1). In the raw gene expression space all models performed similarly (Pearson, Table 1). However, the Pearson correlation values between raw gene expression profiles are strongly influenced by the baseline expression magnitudes of different genes, so we did not consider these metrics meaningful. In the differential expression space (Pearson Delta, Figure 1B, 1C, 1D; Table 1), even the simplest baseline model, “Train Mean” reached better correlation than scGPT, while Random Forest Regressor outperformed scGPT by a large margin. In contrast, using the correlation between the top 20 differentially expressed genes (Pearson Delta DE, Table 1), scGPT had better performance than “Train Mean”. However, in the Top20 DE genes the CRISPR target gene of the perturbation was frequently present. As the target genes’ expression significantly decreased in CRISPRi experiments (Adamson, Replogle) and increased in the CRISPRa experiment (Norman), and scGPT used a perturbation token directly associated with gene tokens, the prediction of target gene expression can be considered trivial. Removing the target genes from the Top20 DE gene list resulted in a decrease in scGPT’s performance in the Top20 DE space (Table 1).

**Table 1.**
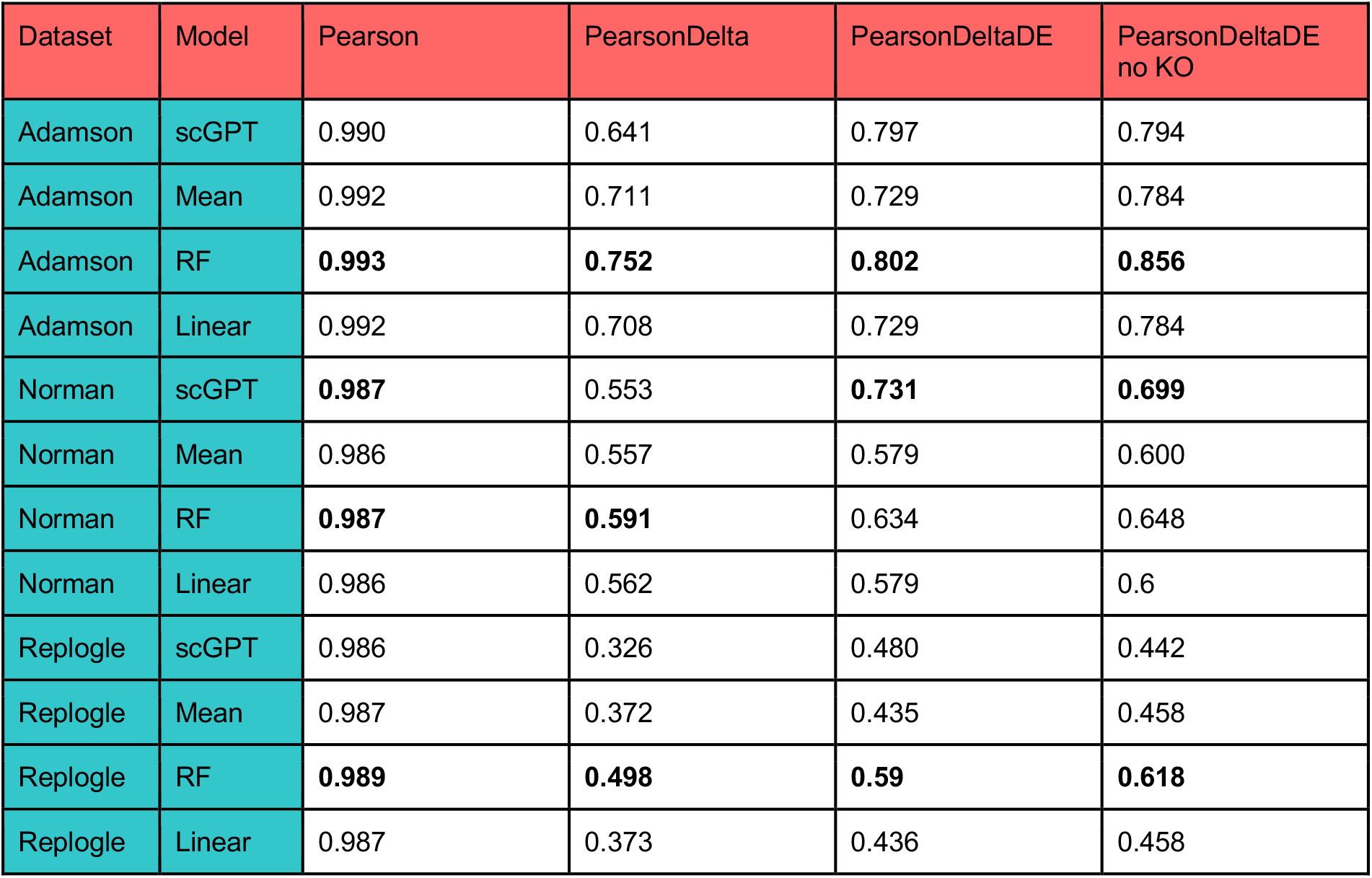
Benchmarking scGPT and baseline models. Evaluation scores for different datasets, models and metrics (detailed in Methods). Best performing scores for each dataset are highlighted with bold.

| Dataset | Model | Pearson | PearsonDelta | PearsonDeltaDE | PearsonDeltaDE no KO |
| --- | --- | --- | --- | --- | --- |
| Adamson | scGPT | 0.990 | 0.641 | 0.797 | 0.794 |
| Adamson | Mean | 0.992 | 0.711 | 0.729 | 0.784 |
| Adamson | RF | **0.993** | **0.752** | **0.802** | **0.856** |
| Adamson | Linear | 0.992 | 0.708 | 0.729 | 0.784 |
| Norman | scGPT | **0.987** | 0.553 | **0.731** | **0.699** |
| Norman | Mean | 0.986 | 0.557 | 0.579 | 0.600 |
| Norman | RF | **0.987** | **0.591** | 0.634 | 0.648 |
| Norman | Linear | 0.986 | 0.562 | 0.579 | 0.6 |
| Replogle | scGPT | 0.986 | 0.326 | 0.480 | 0.442 |
| Replogle | Mean | 0.987 | 0.372 | 0.435 | 0.458 |
| Replogle | RF | **0.989** | **0.498** | **0.59** | **0.618** |
| Replogle | Linear | 0.987 | 0.373 | 0.436 | 0.458 |

In summary we found that even the simplest predictor, the Train Mean model achieved better performance than scGPT regarding predicting the post-treatment RNAseq vectors of perturbed cells, and the simple Random Forest Regressor model outperformed scGPT by a large margin.

## Data composition affects benchmarking results

To explore the unexpected strong performance of the simple Train Mean model, we examined the composition of the benchmark datasets in more detail (Figure 2A). Although the three Perturb-seq datasets contain many single cells (ranging from ∼70,000 in the Adamson dataset to ∼160,000 in the Replogle dataset), the number of distinct perturbations is comparatively much smaller -87 in Adamson, 284 in Norman, and 1,093 in Replogle (Figure 2B). While having multiple single cells per perturbation can be beneficial for model training (i.e., fine-tuning), as it provides insights into perturbation-specific gene expression variability, it would be surprising that large-scale models could efficiently learn and generalise from such a limited number of distinct perturbations.

**Figure 2.**
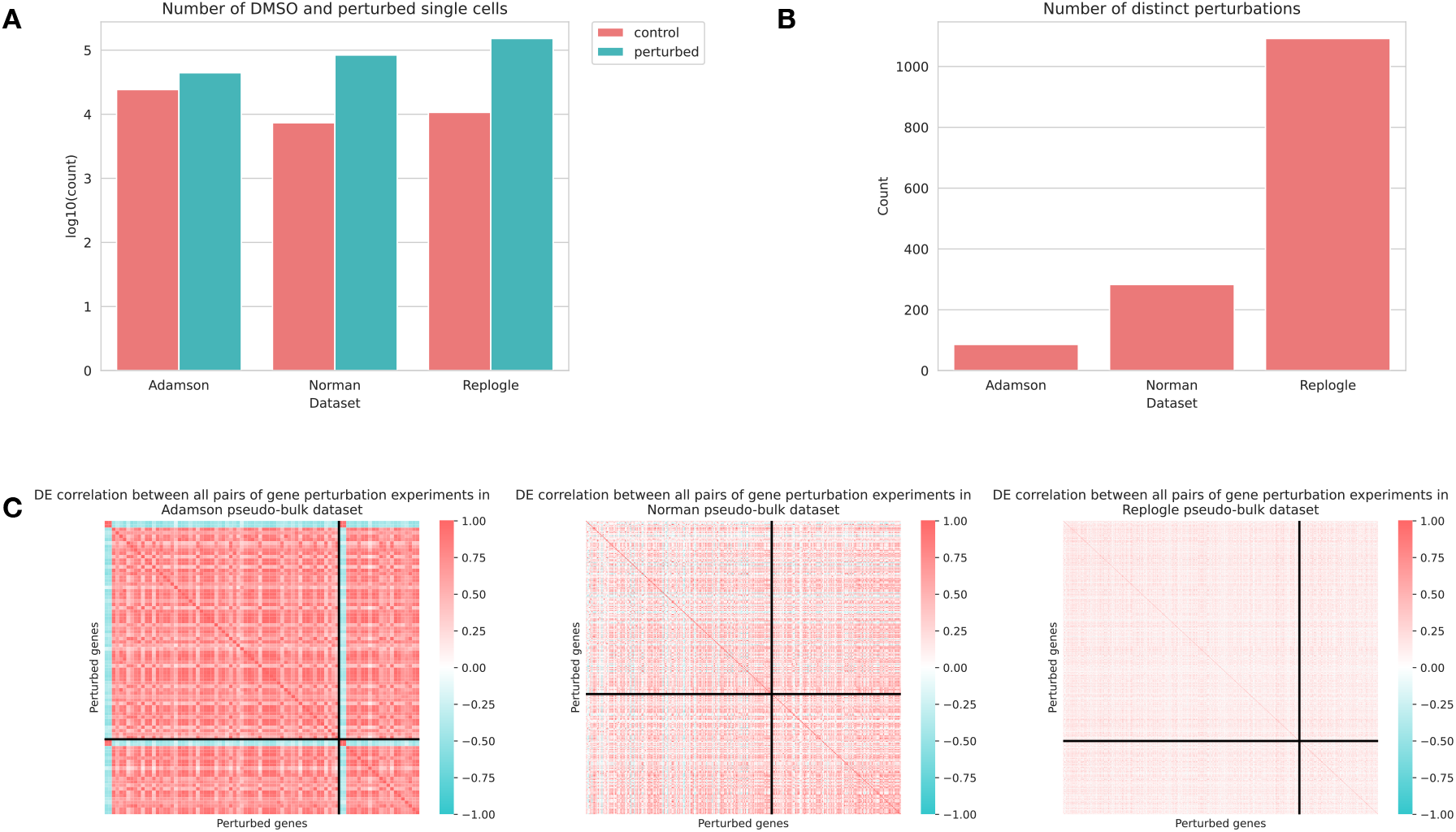
Composition of standard benchmark datasets. (A) Number of single cells (control / perturbed, colour code) in the three benchmark datasets. y axis is log10 scaled (B) Number of distinct perturbations in the three benchmark datasets (C) Correlation heatmaps for pseudo-bulk differential expression signatures for Adamson (left), Norman (middle) and Replogle (right) datasets. The black lines indicate the separation between training and test sets.

To further investigate the benchmark datasets, we analysed the similarities between the pseudo-bulk differential expression profiles across the three datasets (Figure 2C). In the Adamson dataset, we observed high similarity between the perturbation profiles, with a median Pearson correlation of 0.662. This result is not entirely unexpected, given that the Adamson study focused on perturbations specifically targeting endoplasmic reticulum homeostasis, where similar transcriptional responses might be expected. Only a few genes exhibited anti-correlated expression profiles. Similarly, in the Norman dataset, there was a high degree of similarity between perturbation profiles, with a median Pearson correlation of 0.273. In contrast, the Replogle dataset displayed greater variability in the perturbation profiles, with a lower median Pearson correlation of 0.117. This greater variability aligns with the genome-wide scope of the Replogle study, where perturbations target a broader range of biological pathways, leading to more diverse transcriptional outcomes.

In summary, our analysis indicates that while these benchmark datasets contain many single cells, the actual biological variance between the samples is relatively small. This limited variance likely explains why simple models, such as averaging the training examples, perform unexpectedly well in terms of evaluation metrics. However, this small variance also hampers the benchmarks’ ability to effectively differentiate between more complex models, thereby limiting the assessment of their true performance.

## Discussion

In this study, we compared the post-perturbation RNA-seq prediction performance of scGPT against several baseline models using Perturb-seq datasets. Surprisingly, we found that even the simplest baseline, the Train Mean model, outperformed scGPT in most cases. A more advanced machine learning model, the Random Forest Regressor, which incorporated biological prior knowledge in the form of Gene Ontology (GO) terms, outperformed scGPT by a large margin. These results align with a preprint published during the preparation of our manuscript (Ahlmann-Eltze, Huber, and Anders 2024). Our findings also suggest that the current benchmark datasets, particularly Adamson and Norman, may lack sufficient variability to accurately distinguish between the performances of different models.

Given that these benchmark datasets are primarily designed to evaluate the Perturbation Exclusive (PEX) problem - predicting responses to novel perturbations -the correct representation of perturbations is essential for model performance. While scGPT is pre-trained on large-scale, unlabeled single-cell RNA-seq data and likely learns gene regulatory interactions (as shown by the original authors through embedding clustering and pathway analysis), it does not utilise perturbation data during pre-training. Consequently, perturbation-specific information must be learned during fine-tuning via perturbation tokens. The relatively small number of distinct perturbations in the benchmark datasets may not be sufficient to capture this complexity. In contrast, the Random Forest Regressor’s use of GO terms as prior knowledge for perturbation representation appears to provide a more effective means of modelling perturbation responses. This suggests that perturbation effects, which propagate through signalling networks, may be better represented by functional categories such as GO terms than by the gene regulatory networks primarily learned by scGPT.

Our analysis also highlights a significant limitation in the current benchmark datasets - low variance. The Adamson and Norman datasets exhibit high similarity between perturbations, which hinders the ability to distinguish between model performances. Only the Replogle dataset, with its genome-wide scope, demonstrated sufficient variability to provide meaningful benchmarking results. Additionally, the current benchmarks are focused exclusively on the PEX problem, neglecting the Cell Exclusive (CEX) problem, which involves predicting responses in novel cell types. Datasets such as LINCS-L1000 (Subramanian et al. 2017), which feature a larger number of cell lines and perturbations, or Mix-Seq (McFarland et al. 2020), could serve as valuable resources for addressing both PEX and CEX setups and improving future benchmarks.

Our findings also raise questions about the utility of single-cell RNA-seq data for post-perturbation RNA-seq prediction. While single-cell data allow for the modelling of cellular heterogeneity and provide larger sample sizes, the baseline models we used operated on pseudo-bulked data and performed comparably or better than scGPT. Single-cell expression profiles are often subject to technical noise, such as dropout events, which may obscure meaningful biological signals. In our current benchmark study, using single-cell data did not provide an obvious advantage over pseudo-bulked data, particularly given that the datasets were derived from in vitro cell lines with low heterogeneity.

In summary, our benchmarking revealed that scGPT performed comparably to the trivial Train Mean model in post-perturbation RNA-seq prediction tasks and was outperformed by a Random Forest Regressor utilising prior biological knowledge. Furthermore, the low variance in the commonly used benchmark datasets limits their ability to effectively assess model performance. Although our analysis focused on scGPT, other foundational models that have been tested on the same datasets may face similar limitations. While single cell foundational models like scGPT hold promise due to their ability to incorporate large-scale, unlabeled data, several recent studies (Kedzierska et al. 2023; Boiarsky et al. 2023) suggest that their performance on certain tasks may lag behind state-of-the-art or even baseline models. Moving forward, more rigorous and meaningful benchmarks that include higher variance and incorporate diverse datasets are needed to properly assess the applicability of machine learning models in post-perturbation prediction tasks.

## Methods

### scGPT

The scGPT’s results on the GEARS benchmarking datasets were reproduced using the cell-gears v0.0.1 package and a fork of scGPT v0.2.1 (at commit 7301b51).The scGPT fine tunings were performed by the provided tutorial notebook, Tutorial_Perturbation, aligned to each dataset.

### Benchmark datasets

The benchmark datasets were downloaded and processed by the GEARS’ cell-gears v0.0.1 package.

### Model evaluation

The model performance was evaluated using four metrics: 1) Pearson, 2) Pearson Delta, 3) Pearson Delta DE, and 4) Pearson Delta without target genes. The first three metrics were used in the scGPT paper. All metrics were calculated at the ‘bulk’ level, meaning that conditions (control and perturbed states) were mean-aggregated over the gene dimension.

‘Pearson’ refers to the raw correlation between predicted and true post-perturbation expressions. ‘Pearson Delta’ refers to the correlation between the differential expression (post -control) of predicted and true post-perturbation expressions. ‘Pearson Delta DE’ is a variation of Pearson Delta that is calculated only for the top 20 most differentially expressed genes. ‘Pearson Delta without target genes’ excludes the CRISPR target gene(s) from the top 20 most differentially expressed genes.

### Baseline models

scGPT’s performance was compared against three baseline models: 1) Train Mean 2) Random Forest and 3) Elastic Net. The Train Mean model predicts the same vector for each test sample, calculated as the mean post-perturbation expression across all training samples for each perturbation. Random Forest and Elastic Net used embedded GO features for training. PCA dimensionality reduction was applied to the GO matrix (n_genes, n_functions), reducing it to 256 principal components. For each sample, the perturbed gene was identified from the PCA-transformed GO features. In cases where multiple perturbations occurred in a single sample, the embeddings were simply summed. The Random Forest and Elastic Net were trained using the scikit-learn v1.5.2 Python package (Pedregosa et al. 2011). Random forest was trained with the hyperparameter n_estimators=300, while all other hyperparameters were left at their default settings. Elastic Net was trained with alpha=1 and l1_ratio=0.5.

## Acknowledgement

We thank László Mérő and Milán Sztilkovics for the fruitful discussions regarding post-perturbation RNA-seq benchmarking. The authors used ChatGPT to assist with language and grammar of the initial draft manuscript.

## Code and data availability

The code to reproduce our analysis is available at https://github.com/turbine-ai/PerturbSeqPredBenchmark.

## Authors contribution

All authors designed the research and contributed to manuscript writing. GCs performed all analysis.

